# Improving the *COI* DNA barcoding library for Neotropical phlebotomine sand flies (Diptera, Psychodidae)

**DOI:** 10.1101/2022.12.14.520486

**Authors:** Laura Posada-López, Bruno Leite Rodrigues, Ivan Dario Velez, Sandra Uribe

**Affiliations:** PECET (Program for the study and control of tropical diseases), Faculty of Medicine, Universidad de Antioquia, Medellín, Colombia; School of Public Health, University of São Paulo (FSP/USP), São Paulo – SP, Brasil; Grupo de Investigación en Sistemática Molecular, Universidad Nacional de Colombia, Medellín campus, Colombia

**Keywords:** Phlebotominae, single-locus, species delimitation, molecular taxonomy, molecular systematics

## Abstract

A fragment of the mitochondrial *cytochrome c oxidase subunit I* (*COI*) gene was used to generate 156 new barcode sequences for sand flies from different countries of the Neotropical region, mainly Colombia, which had been identified morphologically as 43 species. The sequencing of the *COI* gene allowed the detection of cryptic diversity within species and correctly associated isomorphic females with males identified by morphology. The maximum intraspecific genetic distances ranged from 0 to 8.32% and 0 to 8.92% using uncorrected *p* distances and the K2P model, respectively. The minimum interspecific distance (nearest neighbor) for each species ranged from 1.5 to 14.14% and 1.51 to 15.7% using p and K2P distances, respectively. Three species had more than 3% maximum intraspecific distance: *Psychodopygus panamensis, Micropygomyia cayennensis cayennensis*, and *Pintomyia evansi*. They also were splitted into at least two MOTUs each, using different species delimitation algorithms. Regarding interspecific genetic distances, the species of the genera *Nyssomyia* and *Trichophoromyia* generated values lower than 3% (except *Ny. ylephiletor* and *Ny. trapidoi*). However, the maximum intraspecific distances did not exceed these values, indicating the presence of a barcode gap despite their proximity. Also, nine sand fly species were DNA barcoded for the first time: *Evandromyia georgii, Lutzomyia sherlocki, Ny. ylephiletor, Ny. yuilli pajoti, Psathyromyia punctigeniculata, Sciopemyia preclara, Trichopygomyia triramula, Trichophoromyia howardi*, and *Th. velezbernali*. Thus, the *COI* DNA barcode analysis allowed the correct delimitation of several Neotropical sand fly species from South and Central America and raised questions about the presence of cryptic species for some taxa, which should be further assessed.

## Introduction

The subfamily Phlebotominae Rondani and Berté, in Rondani 1840, comprises about 1,047 species worldwide, 554 in the Neotropical region [1, 2], distributed in 23 genera. Within the subfamily, there are species of particular interest due to their role as vectors of *Leishmania* protozoans, the causative agents of leishmaniasis, which is why their taxonomic determination is crucial to understand biological and ecological aspects that determine patterns and dynamics of the diseases they transmit. Entomological surveillance and species identification are necessary to predict possible risk areas of disease transmission and adopt more efficient control measures [3].

The species-level identification of sand flies is based on morphological characteristics; however, there are limitations, such as phenotypic plasticity within the same species, the presence of cryptic species, isomorphic females, and inappropriate mounting techniques. Additionally, a high degree of skill and taxonomic expertise is required to carry out the identification of some taxa. These aspects point to the need for integrative approaches, which address morphological, molecular, behavioural, and ecological data to better understand the taxonomic status of this subfamily [4, 5].

The DNA barcoding initiative has become an attractive tool in the case of insects of medical importance, where it is necessary to know quickly and accurately which species are present in a transmission area [6-9]. This initiative has been well received due to the connectivity and common language of DNA sequences, which allows researchers from different parts of the world to advance in taxonomic and systematic studies of various groups of organisms, including the vectors of diseases [10, 12].

Some sand fly species have already been processed for the *COI* gene and are available in genetic databases for molecular identification of these taxa (e.g., NCBI GenBank and BOLD Systems). However, there is a knowledge gap in *COI* barcode sequences for some groups, as only a quarter of current species have been sequenced for this marker [12]. Thus, several efforts are being made to make new sequences available, in addition to evaluating their usefulness in the species delimitation within this subfamily [9, 13-19]. Here, we aim to validate the use of *COI* DNA barcoding as a practical tool for species identification, its usefulness for species assignment in isomorphic females, and to evaluate the detection of cryptic diversity that occurs in the same species with different locations.

## Methods

### Sand fly sampling and morphological identification

The sand flies were collected according to the parameters of Colombian decree number 1376 which regulates the permits for specimen collection of biologically diverse wild species for non-commercial research.

The collections were carried out between 2013 and 2016 in nine locations belonging to five Departments of Colombia: Amazonas Department: 1-Leticia (69°56’35’’ S; 4°12’29’’ W) and 2-Puerto Nariño (70°22’59’’ S; 3°46’13’’ W); Antioquia Department : 3-Apartadó (76°37’55’’ S; 7°53’0.9’’ E) and 4-Remedios (74°41’38’’ S; 7°1’39’’ E); Caldas department: 5-Norcasia (74°53’20’’ S; 5°34’27’’ E), 6-Samaná (74°59’34’’ S; 5°24’47’’ E) and 7-Victoria (74°54’45’’ S; 5°18’59’’ E); Magdalena Department: 8-Santa Marta (74°11’56’’ S; 11°14’26’’ E); and Sucre Department: 9-Ovejas (75°13’37’’ S; 9°31’32’’ E). The study locations were selected based on epidemiological studies of leishmaniasis transmission previously carried out by the Program for the Study and Control of Tropical Diseases (PECET). The specimens collected in four Central American countries between 2010 and 2012 were also included; the collections were made as part of a training program carried out by PECET researchers and in association with OPS/OMS (Pan American Health Organization/Organización Panamericana de la Salud). The locations were Nicaragua: 10-Leon/Rota (85° 3’1.7” S; 12°32’53” E); Costa Rica: 11-Limon/San Vicente (84°2’51” S; 9°57’36” E) and 12-Limon/Sibuju (84°2’51” S; 9°57’36” E); Panama: 13-Panama Oeste/Capira-Ollas Arriba (79°54’32” S; 8°48’30” E); and Honduras: 14-Valle/Amapala-El Caracol (87°39’14” S; 13°17’31” E). Figure 1 shows a map of the 14 locations. The collections carried out on private property received verbal permission from the landowners before the sampling.

**Figure 1.**
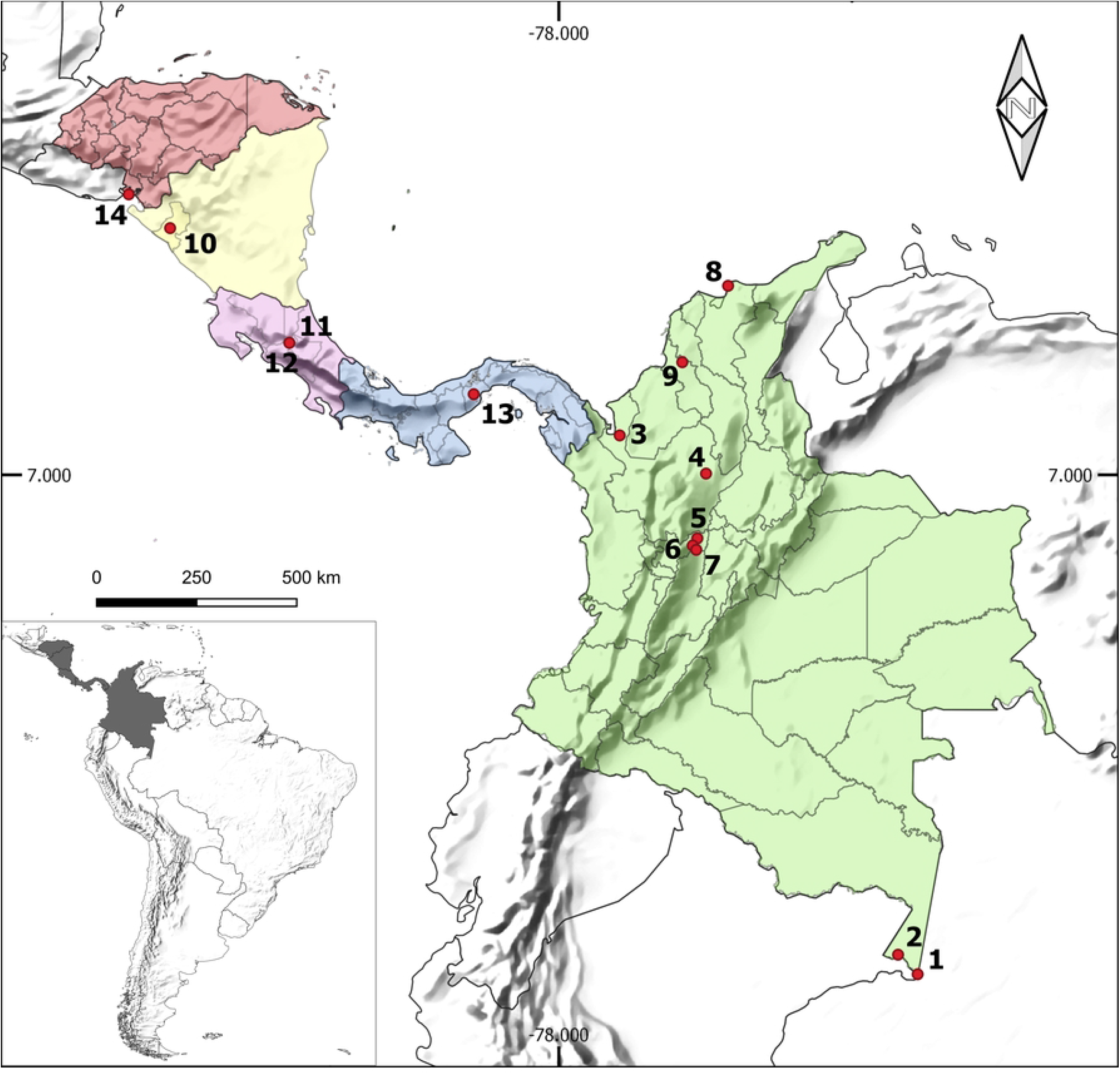
Map showing the sampling sites of sand flies species used in this study. Colombia: 1-Leticia (Amazonas), 2-Puerto Nariño (Amazonas), 3-Apartadó (Antioquia) 4-Remedios (Antioquia), 5-Norcasia (Caldas), 6-Samaná (Caldas), 7-Victoria (Caldas), 8-Santa Marta (Magdalena), 9-Ovejas (Sucre). Nicaragua: 10-Leon/Rota, Costa Rica: 11-Limon/San Vicente, 12-Limon/Sibuju, Panamá: 13-Panamá Oeste/Capira-Ollas Arriba and Honduras: 14-Valle/Amapala-El Caracol.

Sand flies were collected using CDC light traps located in peridomiciliary environments, preferably near domestic animal shelters, where insects are more abundant, and forested fragments.

The insects were stored dry and subsequently processed in the Medical and Molecular Entomology Laboratory of PECET at the University of Antioquia. The thorax was stored dry at −20° C until it was processed by molecular techniques. The head and abdomen were used for the morphological identification following Galati (2013) [1], and the Marcondes nomenclature (2007) [20] was followed for the abbreviations of the genera.

### DNA extraction, PCR and sequencing

Total DNA from each specimen was extracted from the remaining parts of sand flies (thorax) using the high salt concentration protocol described by Porter and Collins (1991) [21]. The *COI* gene fragment was amplified using the primers: LCO1490 (5′ GGT CAA CAA ATC ATA AAG ATA TTG G 3′) and HCO2198 (5′ TAA ACT TCA GGG TGA CCA AAA AAT CA 3′) [22]. The PCR products were visualized on electrophoresis using 1% agarose gel and sequenced in both chain directions by Macrogen Inc. (Korea).

### Sequence Analysis

The obtained chromatograms were edited using the BioEdit v7.0.9 software [23] to generate a consensus sequence for each specimen. All sequences were aligned using the ClustalW algorithm and then visually examined to ensure there were no stop codons, pseudogenes, and nuclear copies of mitochondrial origin (NUMTs) using the Mega v7 software [24]. The *COI* sequences were then submitted to the BOLD Systems database [25] and are available in the “CLBAR – Improving the DNA barcoding library for Neotropical sand flies” project and NCBI GenBank [26] database, being assigned the Accession Numbers OP964207 - OP964362.

The sequence alignment was done using MUSCLE [27] implemented in MEGA v7. Pairwise genetic distances for both maximum intraspecific and minimum interspecific (nearest-neighbor, NN) were generated in the BOLD Systems environment using the “Barcode Gap Analysis” tool with uncorrected (p distances) or the Kimura-2-parameter (K2P) models. The consensus alignment was then used to generate a phenogram using the neighbor-joining (NJ) method with pairwise genetic distances and 1,000 bootstrap pseudoreplicates in the software MEGA v7. Also, a phylogenetic gene tree was generated using the maximum likelihood (ML) method in the software RAxML v8 [28] and its graphical user interface, raxmlGUI v2.0 [29]. For the ML tree, the GTR+G+I substitution model was used as suggested by jModelTest v2 [30], and the data were partitioned according to codon position. A sequence of Sycorax konopiki (KT946601.1) was included as an outgroup to root the ML tree.

The DNA barcode sequences were also identified at the molecular operational taxonomic unit (MOTU) level, which are groups of specimens based on their molecular similarity at a given molecular marker [31]. Several algorithms were designed to sort barcode sequences of a given dataset into MOTUs without a priori information (i.e., discovery approaches [32]). Therefore, several single-locus species delimitation methods were used to associate morphologically distinct species with MOTUs, evaluating the usefulness of *COI* DNA barcodes for the taxonomy of different sand flies from the Neotropical region. For this, the following methods were employed: i) Automatic Barcode Gap Discovery (ABGD) [33]; ii) Refined Single Linkage (RESL) [34]; iii) TCS haplotype networks using statistical parsimony method [35-36]; and iv) Poisson Tree Processes (PTP) [37]. The ABGD analysis (available at https://bioinfo.mnhn.fr/abi/public/abgd/abgdweb.html) clusters sequences according to their similarity in a given genetic pairwise distance matrix according to inferred barcode gaps in the dataset and then recursively applies this procedure to the obtained MOTUs to get finer partitions. Two different ABGD analyses were run using uncorrected p distances and the K2P model, and the parameters Pmin=0.005, Pmax=0.1, and X=1.0. For ABGD, it was considered the recursive partitions generated with a range of prior intraspecific divergence between 1% and 2.5% [16]. The RESL algorithm was designed to deal with large amounts of DNA barcode sequences in the BOLD Systems (https://boldsystems.org/) and operates by linking similar sequences and then optimizing the MOTU delimitation with a graphic analytical approach using a Markov Clustering (MCL). RESL analysis was run inside the BOLD environment using the ‘cluster sequences’ tool and default parameter. The software TCS v1.21 infers haplotype networks using the statistical parsimony method and can be used for species delimitation. This approach can generate disconnected networks for different morphospecies while analyzing sand fly-DNA barcode datasets [38-39]. The networks inferred by TCS were visualized and edited using the tcsBU web server [40] (available at https://cibio.up.pt/software/tcsBU/). Lastly, the algorithm PTP is a coalescent-derived method that seeks to differentiate stochastic population processes from speciation events in a phylogenetic gene tree. Therefore, the MOTU delimitation by PTP was conducted by submitting the ML gene tree (after pruning the outgroup) to the web server (available at https://species.h-its.org/ptp/) and using its default settings, except for the number of MCMC generations, which was changed to 500,000.

## Results

One hundred fifty-six new *COI* barcode sequences were generated for phlebotomine sand flies from different countries of the Neotropical region, mainly from Colombia (Table 1). The morphological identification of these specimens revealed the presence of 43 species, of which 41 were identified at the species level while the other two were assigned only at subgenus, *Lutzomyia* (*Tricholateralis*) Galati, 2003 and series level *Psychodopygus* Guyanensis series Barretto, 1962. Regarding the species identification, the taxa belong to the following genera: *Brumptomyia, Evandromyia, Lutzomyia*, Micropygomyia, *Nyssomyia, Pintomyia, Pressatia, Psathyromyia, Psychodopygus, Sciopemyia, Trichophoromyia*, and *Trichopygomyia*. The complete information on the analyzed species and sample locations is listed in Tables 1 and S1.

**Table 1.**
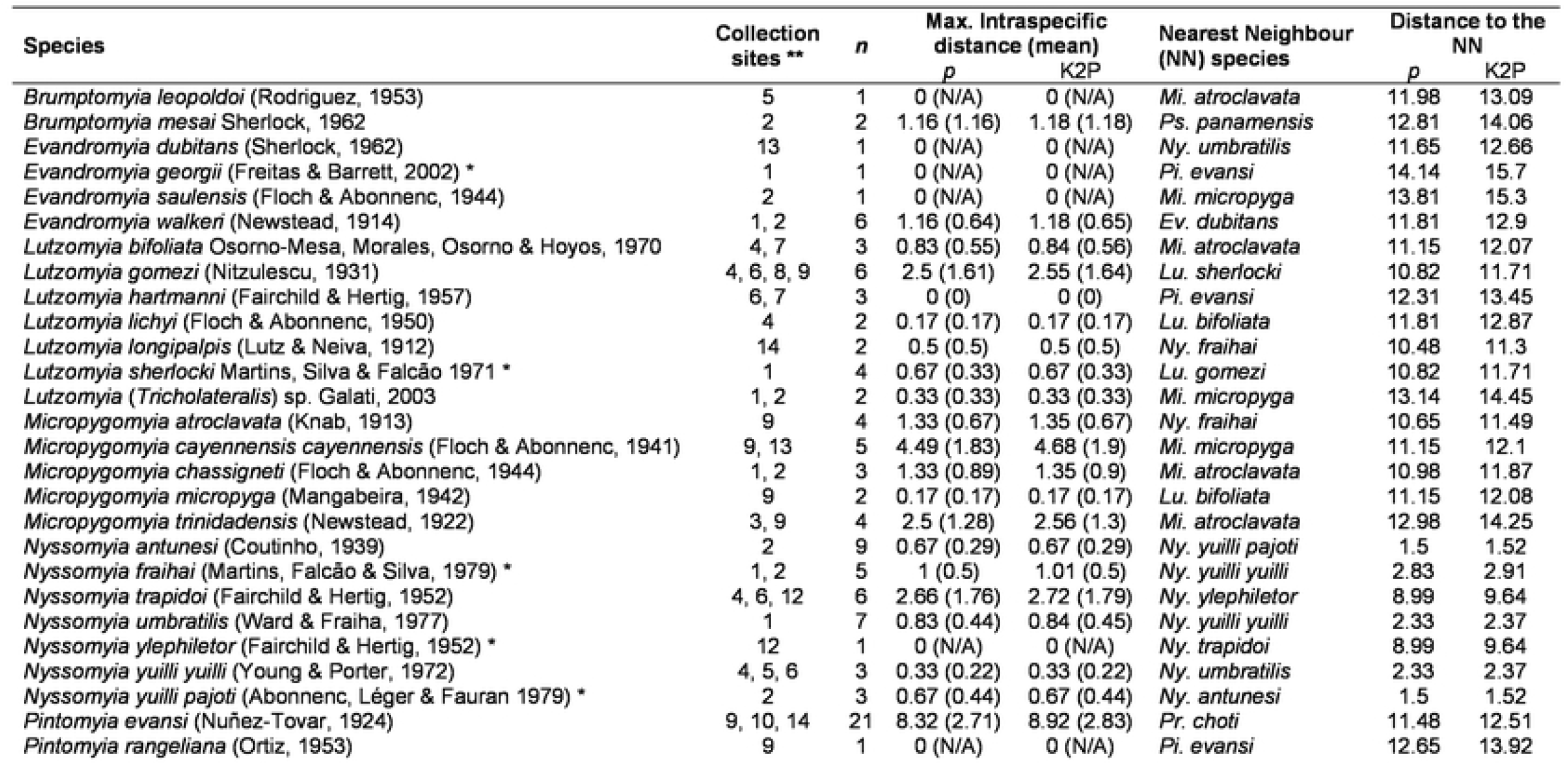
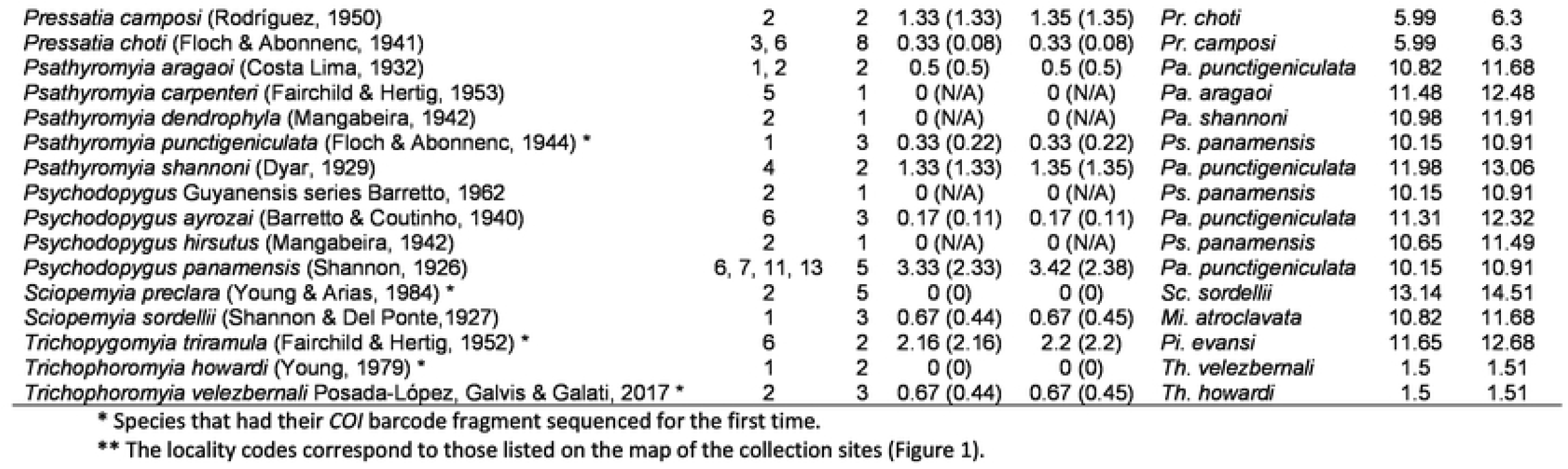
Nominal species, collection sites, maximum intraspecific genetic divergence, and the minimum distance to the nearest neighbour of sand fly species from the Neotropical region analysed in this study.

| Species | Collection sites ** | n | Max. Intraspecific distance (mean) |  | Nearest Neighbour (NN) species | Distance to the NN |  |
| --- | --- | --- | --- | --- | --- | --- | --- |
|  |  |  | p | K2P |  | p | K2P |
| *Brumptomyia leopoldoi* (Rodríguez, 1953) | 5 | 1 | 0 (N/A) | 0 (N/A) | *Mi. atroclavata* | 11.98 | 13.09 |
| *Brumptomyia mesai* Sherlock, 1962 | 2 | 2 | 1.16 (1.16) | 1.18 (1.18) | *Ps. panamensis* | 12.81 | 14.06 |
| *Evandromyia dubitans* (Sherlock, 1962) | 13 | 1 | 0 (N/A) | 0 (N/A) | *Ny. umbratilis* | 11.65 | 12.66 |
| *Evandromyia georgii* (Freitas & Barrett, 2002) * | 1 | 1 | 0 (N/A) | 0 (N/A) | *Pi. evansi* | 14.14 | 15.7 |
| *Evandromyia saulensis* (Floch & Abonnenc, 1944) | 2 | 1 | 0 (N/A) | 0 (N/A) | *Mi. micropyga* | 13.81 | 15.3 |
| *Evandromyia walkeri* (Newstead, 1914) | 1, 2 | 6 | 1.16 (0.64) | 1.18 (0.65) | *Ev. dubitans* | 11.81 | 12.9 |
| *Lutzomyia bifoliata* Osorno-Mesa, Morales, Osorno & Hoyos, 1970 | 4, 7 | 3 | 0.83 (0.55) | 0.84 (0.56) | *Mi. atroclavata* | 11.15 | 12.07 |
| *Lutzomyia gomezi* (Nitzulescu, 1931) | 4, 6, 8, 9 | 6 | 2.5 (1.61) | 2.55 (1.64) | *Lu. sherlocki* | 10.82 | 11.71 |
| *Lutzomyia hartmanni* (Fairchild & Hertig, 1957) | 6, 7 | 3 | 0 (0) | 0 (0) | *Pi. evansi* | 12.31 | 13.45 |
| *Lutzomyia lichyi* (Floch & Abonnenc, 1950) | 4 | 2 | 0.17 (0.17) | 0.17 (0.17) | *Lu. bifoliata* | 11.81 | 12.87 |
| *Lutzomyia longipalpis* (Lutz & Neiva, 1912) | 14 | 2 | 0.5 (0.5) | 0.5 (0.5) | *Ny. fraihai* | 10.48 | 11.3 |
| *Lutzomyia sherlocki* Martins, Silva & Falcão 1971 * | 1 | 4 | 0.67 (0.33) | 0.67 (0.33) | *Lu. gomezi* | 10.82 | 11.71 |
| *Lutzomyia* (*Tricholateralis*) sp. Galati, 2003 | 1, 2 | 2 | 0.33 (0.33) | 0.33 (0.33) | *Mi. micropyga* | 13.14 | 14.45 |
| *Micropygomyia atroclavata* (Knab, 1913) | 9 | 4 | 1.33 (0.67) | 1.35 (0.67) | *Ny. fraihai* | 10.65 | 11.49 |
| *Micropygomyia cayennensis cayennensis* (Floch & Abonnenc, 1941) | 9, 13 | 5 | 4.49 (1.83) | 4.68 (1.9) | *Mi. micropyga* | 11.15 | 12.1 |
| *Micropygomyia chassigneti* (Floch & Abonnenc, 1944) | 1, 2 | 3 | 1.33 (0.89) | 1.35 (0.9) | *Mi. atroclavata* | 10.98 | 11.87 |
| *Micropygomyia micropyga* (Mangabeira, 1942) | 9 | 2 | 0.17 (0.17) | 0.17 (0.17) | *Lu. bifoliata* | 11.15 | 12.08 |
| *Micropygomyia trinidadensis* (Newstead, 1922) | 3, 9 | 4 | 2.5 (1.28) | 2.56 (1.3) | *Mi. atroclavata* | 12.98 | 14.25 |
| *Nyssomyia antunesi* (Coutinho, 1939) | 2 | 9 | 0.67 (0.29) | 0.67 (0.29) | *Ny. yuilli pajoti* | 1.5 | 1.52 |
| *Nyssomyia fraihai* (Martins, Falcão & Silva, 1979) * | 1, 2 | 5 | 1 (0.5) | 1.01 (0.5) | *Ny. yuilli yuilli* | 2.83 | 2.91 |
| *Nyssomyia trapidoi* (Fairchild & Hertig, 1952) | 4, 6, 12 | 6 | 2.66 (1.76) | 2.72 (1.79) | *Ny. ylephiletor* | 8.99 | 9.64 |
| *Nyssomyia umbratilis* (Ward & Fraiha, 1977) | 1 | 7 | 0.83 (0.44) | 0.84 (0.45) | *Ny. yuilli yuilli* | 2.33 | 2.37 |
| *Nyssomyia ylephiletor* (Fairchild & Hertig, 1952) * | 12 | 1 | 0 (N/A) | 0 (N/A) | *Ny. trapidoi* | 8.99 | 9.64 |
| *Nyssomyia yuilli yuilli* (Young & Porter, 1972) | 4, 5, 6 | 3 | 0.33 (0.22) | 0.33 (0.22) | *Ny. umbratilis* | 2.33 | 2.37 |
| *Nyssomyia yuilli pajoti* (Abonnenc, Léger & Fauran 1979) * | 2 | 3 | 0.67 (0.44) | 0.67 (0.44) | *Ny. antunesi* | 1.5 | 1.52 |
| *Pintomyia evansi* (Nuñez-Tovar, 1924) | 9, 10, 14 | 21 | 8.32 (2.71) | 8.92 (2.83) | *Pr. choti* | 11.48 | 12.51 |
| *Pintomyia rangeliana* (Ortiz, 1953) | 9 | 1 | 0 (N/A) | 0 (N/A) | *Pi. evansi* | 12.65 | 13.92 |
| *Pressatia camposi* (Rodríguez, 1950) | 2 | 2 | 1.33 (1.33) | 1.35 (1.35) | *Pr. choti* | 5.99 | 6.3 |
| *Pressatia choti* (Floch & Abonnenc, 1941) | 3, 6 | 8 | 0.33 (0.08) | 0.33 (0.08) | *Pr. camposi* | 5.99 | 6.3 |
| *Psathyromyia aragaoi* (Costa Lima, 1932) | 1, 2 | 2 | 0.5 (0.5) | 0.5 (0.5) | *Pa. punctigeniculata* | 10.82 | 11.68 |
| *Psathyromyia carpenteri* (Fairchild & Hertig, 1953) | 5 | 1 | 0 (N/A) | 0 (N/A) | *Pa. aragaoi* | 11.48 | 12.48 |
| *Psathyromyia dendrophyla* (Mangabeira, 1942) | 2 | 1 | 0 (N/A) | 0 (N/A) | *Pa. shannoni* | 10.98 | 11.91 |
| *Psathyromyia punctigeniculata* (Floch & Abonnenc, 1944) * | 1 | 3 | 0.33 (0.22) | 0.33 (0.22) | *Ps. panamensis* | 10.15 | 10.91 |
| *Psathyromyia shannoni* (Dyar, 1929) | 4 | 2 | 1.33 (1.33) | 1.35 (1.35) | *Pa. punctigeniculata* | 11.98 | 13.06 |
| *Psychodopygus Guyanensis* series Barretto, 1962 | 2 | 1 | 0 (N/A) | 0 (N/A) | *Ps. panamensis* | 10.15 | 10.91 |
| *Psychodopygus ayrozai* (Barretto & Coutinho, 1940) | 6 | 3 | 0.17 (0.11) | 0.17 (0.11) | *Pa. punctigeniculata* | 11.31 | 12.32 |
| *Psychodopygus hirsutus* (Mangabeira, 1942) | 2 | 1 | 0 (N/A) | 0 (N/A) | *Ps. panamensis* | 10.65 | 11.49 |
| *Psychodopygus panamensis* (Shannon, 1926) | 6, 7, 11, 13 | 5 | 3.33 (2.33) | 3.42 (2.38) | *Pa. punctigeniculata* | 10.15 | 10.91 |
| *Sciopemyia preclara* (Young & Arias, 1984) * | 2 | 5 | 0 (0) | 0 (0) | *Sc. sordellii* | 13.14 | 14.51 |
| *Sciopemyia sordellii* (Shannon & Del Ponte, 1927) | 1 | 3 | 0.67 (0.44) | 0.67 (0.45) | *Mi. atroclavata* | 10.82 | 11.68 |
| *Trichopygomyia triramula* (Fairchild & Hertig, 1952) * | 6 | 2 | 2.16 (2.16) | 2.2 (2.2) | *Pi. evansi* | 11.65 | 12.68 |
| *Trichophoromyia howardi* (Young, 1979) * | 1 | 2 | 0 (0) | 0 (0) | *Th. velezbernali* | 1.5 | 1.51 |
| *Trichophoromyia velezbernali* Posada-López, Galvis & Galati, 2017 * | 2 | 3 | 0.67 (0.44) | 0.67 (0.45) | *Th. howardi* | 1.5 | 1.51 |

\* Species that had their *COI* barcode fragment sequenced for the first time.
\*\* The locality codes correspond to those listed on the map of the collection sites (Figure 1).

**Table S1.** Sample IDs, BOLD processes IDs, GenBank accession numbers, nominal species, sex of the specimens, and collection sites of sand fly species from the Neotropical region analysed in this study.

The sequencing resulted in a consensus alignment of 601 bp of the standard *COI* DNA barcode fragment described by Folmer (1994) [22]. The visual inspection of the alignment indicates the absence of stop codons in the middle of sequences, pseudogenes and/or nuclear copies of mitochondrial origin (NUMT).

The number of barcoded specimens per species ranged from one to 21. The maximum intraspecific genetic distances ranged from 0 to 8.32% and 0 to 8.92% using uncorrected p distances and the K2P model, respectively (Table 1). The minimum interspecific distance (Nearest Neighbor) for each species ranged from 1.5 to 14.14% and 1.51 to 15.7% using p and K2P distances, respectively (Table 1). Three species had more than 3% of maximum intraspecific distance: *Psychodopygus* panamensis (3.33% using p distances; 3.42% with the K2P model), *Micropygomyia cayennensis cayennensis* (4.49; 4.68), and *Pintomyia evansi* (8.32; 8.92), but their distances to the nearest neighbors were 10.15/10.91, 11.15/12.1, and 11.48/12.51, respectively (Table 1). Regarding interspecific genetic distances, the species of the genera *Nyssomyia* and *Trichophoromyia* showed values lower than 3% (except for *Ny. ylephiletor* and *Ny. trapidoi*) for both p and K2P distances (Table 1). However, the maximum intraspecific distances did not exceed these values, indicating the presence of a barcode gap despite their proximity.

The NJ phenogram and ML phylogenetic tree grouped conspecific sequences into well-supported clusters/clades for all the analyzed species, sometimes splitting them into more than one group (Figures 2 and S1). Some close-related species of the genera *Nyssomyia* and *Trichophoromyia* formed different clusters comprising only one nominal species each. However, the association of species-level identified males with females of the two *Trichophoromyia* species was somehow hampered due to the lack of diagnostic characters (Figure 3). All analyses separated the species *Ny. yuilli yuilli* from *Ny. fraihai*, the latter only collected in the Colombian Amazon. These results suggest that it is very likely that they are two different species; however, the separation by morphology was not possible. Further, the ML tree indicates that the species pair *Ny. yuilli pajoti*/*Ny. antunesi* may show a paraphyletic pattern (Figure S1). On the other hand, NJ analysis splitted *Ps. panamensis, Mi. trinidadensis, Pi. evansi, Mi. cayennensis cayennensis*, and *Lu. gomezi* into at least two well-supported clades which agree with the samples’ geographical location, except *Lu. gomezi* (Figure 2).

**Figure 2.**
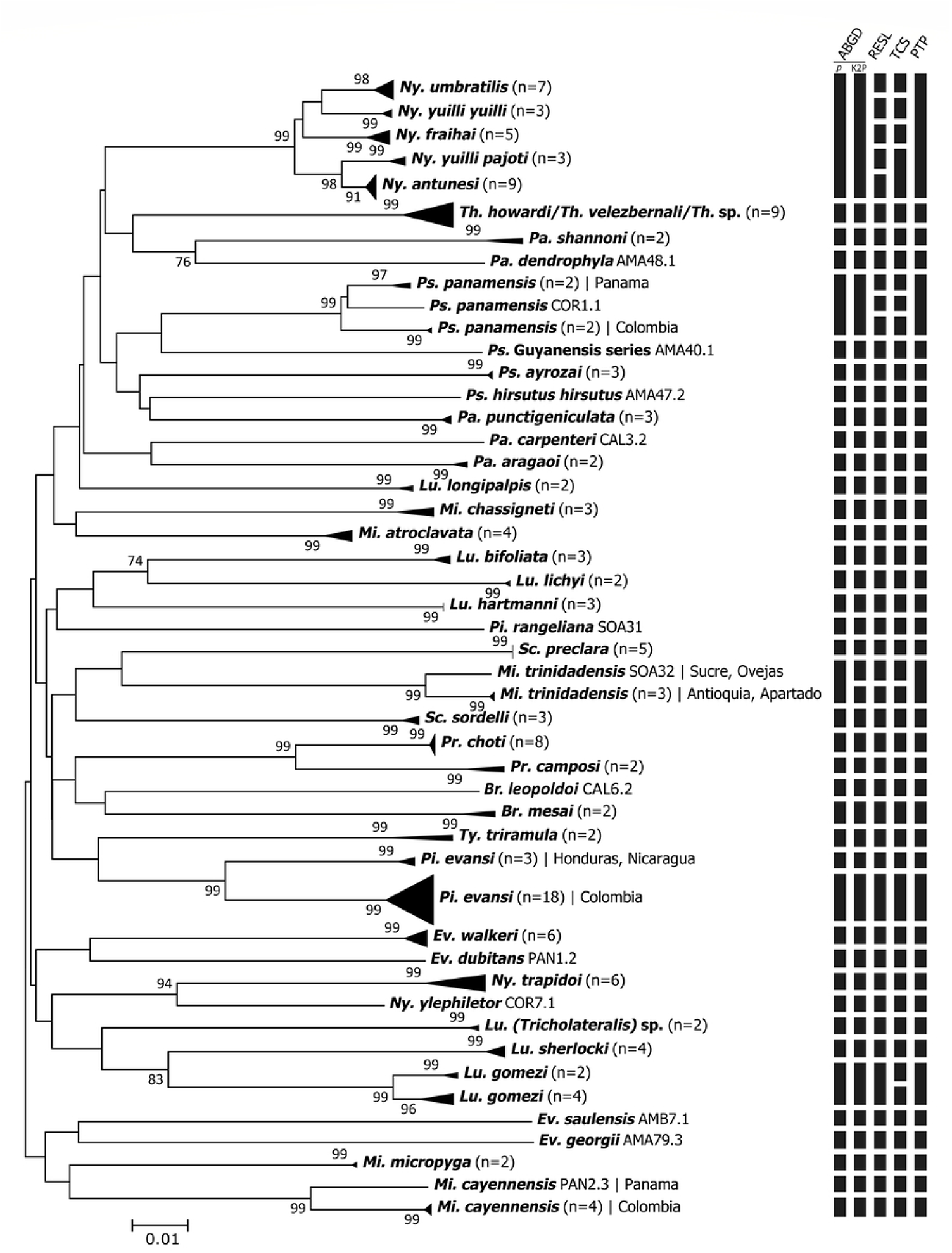
Neighbor-joining phenogram of *COI* sequences of sand fly species from the Neotropical Region. Numbers near nodes indicate bootstrap values above 70. Lateral black bars indicate the MOTU species delimitation partitions made by the algorithms ABGD, RESL, TCS, and PTP.

**Figure 3.**
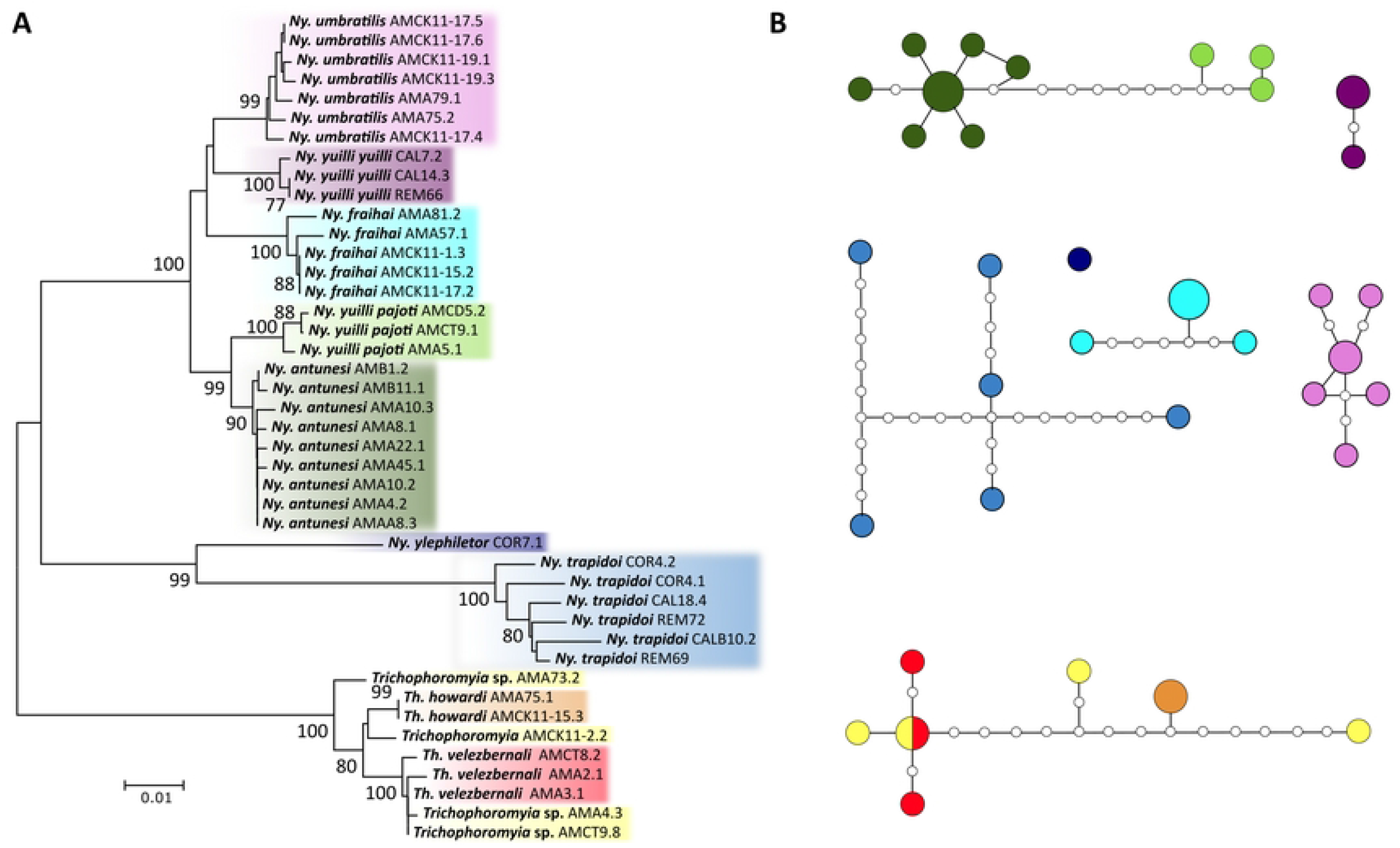
(A) Neighbor-joining phenogram of *COI* sequences of the sand fly genera *Nyssomyia* and *Trichophoromyia*. Numbers near nodes indicate bootstrap values above 70. Clusters are colored according to the nominal species. (B) TCS haplotype network analysis of *COI* sequences of the sand fly genera *Nyssomyia* and *Trichophoromyia*. Unconnected networks are delimited MOTUs and are colored according to nominal species and NJ patterns. Each circle represents a unique haplotype, with its size proportional to the number of individuals, and the small white circles represent inferred unobserved haplotypes.

**Figure S1**. Phylogenetic gene tree based on *COI* DNA barcode sequences of Neotropical sand flies. Numbers near nodes indicate bootstrap values above 70, except the clade comprising Psychodopygina species, which is highlighted in bold.

Regarding species delimitation analyses, the algorithms ABGD (using p and K2P distances), RESL, TCS, and PTP clustered the barcode sequences of the 43 morphospecies into 40, 41, 47, 47, and 40 MOTUs, respectively (Figure 2). ABGD and PTP generated the most conservative partition since, in both cases, five of the nominal species within the *Nyssomyia* genus merged into the same MOTU, and the same happened for *Th. howardi* and *Th. velezbernali* (Figure 2). The algorithms RESL and TCS correctly partitioned the barcode sequences into MOTUs according to the morphospecies but also merged the pairs *Ny. yuilli pajoti*/*Ny. antunesi* (only for TCS), and *Th. howardi*/*Th. velezbernali* (Figure 2). Differently, some or all algorithms splitted *Ps. panamensis, Mi. trinidadensis, Pi. evansi, Mi. cayennensis cayennensis*, and *Lu. gomezi* into at least two well-supported clades which agree with the well-supported clusters/clades of the NJ and ML analysis (Figures 2 and S1).

## Discussion

This study helps improve the digital repository of barcode sequences, the Barcode of Life Data Systems – BOLD, which was designed to assemble and organize all barcode sequence records and provide tools for analyzing these sequences [25]. New *COI* barcode sequences were generated for sand fly species that had not been previously processed. Furthermore, our results indicate that *COI*-DNA barcoding is a useful tool to delimit and identify different sand fly species from the Neotropical region. Different single-locus species delimitation methods were employed to analyze nucleotide divergences from different perspectives, and in almost all cases, the nominal species were assigned as belonging to at least one MOTU.

This study generated the first *COI* sequences for nine sand fly species: *Evandromyia georgii, Lutzomyia sherlocki, Ny. ylephiletor, Ny. yuilli pajoti, Psathyromyia punctigeniculata, Sciopemyia preclara, Trichopygomyia triramula, Trichophoromyia howardi*, and *Th. velezbernali*. All these cases were compared to related species within the same genus, and all seemed to have unique barcode sequences – except for some cases of *Nyssomyia* and *Trichophoromyia* genera – which can be used for future molecular identification of these taxa. Some other studies have evaluated the usefulness of *COI* barcodes in the identification of sand flies from Central America [7, 41] and Colombia [9, 17, 42-43]. Moreover, in Colombia, the sampling efforts were carried out mainly in the Caribbean and Andean regions of the country, which can have different sand fly fauna compared with the southeast region [44]. Consequently, our study surveyed sand flies from the Amazon region of Colombia, in the municipalities of Puerto Nariño and Leticia, close to the borders of Peru and Brazil. Most of the new DNA barcode records comprise specimens from these locations (Table 1, Figure 1).

For this new dataset, various algorithms were employed for species delimitation to associate morphologically distinct species with MOTUs. In general, both the NJ/ML analysis and the species delimitation made by ABGD/RESL/TCS/PTP provide consistent results concerning the nominal species (Figure 2). However, the distance and tree-based methods – ABGD and PTP – were more conservative and merged some close-related species, while RESL and TCS worked well for our dataset, providing more reliable partitions with the sampled species. None of these methods on their own are really capable of accurately delimiting evolutionary lineages due to their limitations when analyzing single-locus DNA barcode data without a priori information on the species boundaries, therefore, it is preferable to use a set of these methods and other lines of evidence to propose putative species [32]. The algorithms used here see the data in different ways, so the congruence between them may indicate that the resulting delimitation is probably correct, but the disagreements should be analyzed in a parsimony way [45]. Although some algorithms merged some species within the *Nyssomyia* and *Trichophoromyia* genera, it does not mean that the delimitation of these taxa by morphology is incorrect.

The pairwise genetic distances – whether uncorrected or using the K2P model – indicate the presence of a ‘barcode gap’ within species and their nearest neighbor (Table 1). This pattern is usually used to define the *COI* barcode as an excellent molecular marker for species identification [46], including sand flies [9, 15], but the clear distinction between these two classes of distances (intra and interspecific) may overlap while analyzing larger datasets with close-related taxa [16, 19, 39, 47-48]. Indeed, it is impossible to establish a generalized standard limit between intra and inter-species barcode divergence to many groups of organisms. This seems to be true even when analyzing species within the same sand fly subgenus (e.g., *Evandromyia (Aldamyia)*, [39]). The extent of the overlap and the absence of the so-called ‘barcode gap’ should not interfere with the rate of successful identifications by DNA barcodes of insects [48].

Regarding the closely related species *Ny. yuilli yuilli* and *Ny. fraihai*, the pairwise genetic distance, and NJ analysis indicate the separation into two clusters. These two nominal species have isomorphic females, but *Ny. yuilli yuilli* is restricted to the Andean and trans-Andean regions of Colombia, while *Ny. fraihai* is widely distributed in Brazil’s Amazon and Atlantic Forest regions [49]. Here, we correctly associated the isomorphic females of the Andean region with males morphologically-identified as *Ny. yuilli yuilli*, but the same was not possible regarding Amazonian specimens since it was only possible to collect females in these locations. However, these Amazonian females were assigned as *Ny. fraihai*, as the collection sites border Brazilian states located in the Amazon region, in which this species has already been reported [50]. A unique *COI* barcode sequence of *Ny. fraihai* (GenBank accession: KP112771) was previously generated from a male specimen collected near a type location in the state of Bahia, in the Atlantic Forest region of Brazil [16, 49], but the nucleotide distance analyses indicated that this individual could represent a different species from those analyzed in this study (data not shown). Therefore, future studies that analyze a greater number of *Ny. fraihai* specimens of different sexes from both biomes, Atlantic Forest and Amazon, should assess the presence of possible new species of these taxa.

Here, the taxa *Ny. antunesi, Ny. fraihai, Ny. umbratilis, Ny. yuilli yuilli, Ny. yuilli pajoti, Th. howardi*, and *Th. velezbernali* did not reach even 3% of minimum divergence to the NN, which is usually seen in other sand fly species as an intraspecific distance value (Table 1). Nevertheless, despite the low values, it did not prevent the formation of individual genetic clusters for all nominal species in the NJ analysis (Figure 3). However, some species delimitation algorithms merged the above-mentioned *Nyssomyia* and *Trichophoromyia* species into a single MOTU each, and for the species pair *Ny. antunesi*/*Ny. yuilli pajoti*, it was not possible to form monophyletic clades in the ML gene tree (Figure S1). According to the phylogenetic systematic analysis of morphological data, these two genera belong to the Psychodopygina subtribe and are considered the most derived groups [50]. Other studies that analyzed *COI* barcodes of *Nyssomyia* spp. from Brazil indicate that the nucleotide divergence between species in this genus is low and may differ from other sand fly taxa [14, 16, 19]. Despite this, *Ny. trapidoi* and *Ny. ylephiletor* achieved a reasonable degree of interspecific genetic divergence, being correctly delimited in all species delimitation algorithms.

Regarding the molecular taxonomy status of the genus *Trichophoromyia*, little information is found in the literature, and *COI* barcode sequences are publicly available for only three species: *Th. reburra, Th. ininii, and Th. viannamartinsi*, which were sequenced and analyzed in different studies [9, 16]. Considering the incredible richness of this genus and the fact that most females are isomorphic [50], this knowledge gap must be filled. In the present study, more than one species of *Trichophoromyia* were analyzed for the first time, which appear to have similar, or even smaller, nucleotide distances than those of the genus *Nyssomyia* (Table 1). In fact, all species delimitation algorithms merged *Th. howardi* and *Th. velezbernali* into a single MOTU, but the NJ and TCS analysis indicates the absence of shared haplotypes, at least for male specimens (Figure 3). Further, some female specimens – which are isomorphic for these two species – could not be correctly associated with males due to a lack of informative characters (Figure 3). Our sampling effort was not satisfactory for this taxa, and future studies may elucidate the actual taxonomic status of these and other species of *Trichophoromyia* using multilocus efforts, which are highly recommended when there are poly and paraphyletic patterns in the genealogy of the alleles studied due to recent speciation processes [51].

One of the main benefits of integrating molecular data into insect taxonomy is the association of immature life stages of holometabolous taxa with adults and isomorphic females with males identified by morphology [52-53]. The present study focused only on generating *COI* sequences for adult specimens. Some species of our dataset have females that are indistinguishable using morphological characters, and it was only possible to correctly associate male and female specimens of the taxa *Mi. cayennensis cayennensis, Mi. chassigneti, Lu. hartmanni, Pr. choti, Pr. camposi, Ty. triramula*. These findings are increasingly relevant because the entomological monitoring of sand flies and leishmaniases in endemic areas is based on species-level identification, especially female specimens, which actively participate in the transmission of pathogens. Highlighting the usefulness of *COI* barcodes in identifying these species may contribute to the use of this tool for monitoring these insects, which can also help identify vector species in studies that assess vector competence/capacity and natural infection by *Leishmania*. In addition, this correct association indicates that the morphological identification of vouchers can be examined in more detail to assess the existence of other morphological characteristics for identifying the sexes that are considered isomorphic until then. Other sand fly DNA barcoding efforts established this molecular marker as relevant for this type of association in the genera *Brumptomyia* [16], *Psychodopygus* [19], and *Phlebotomus* [38]. Therefore, the sequencing of complex groups in which several females are isomorphic, such as the genera *Brumptomyia, Lutzomyia, Pintomyia Townsendi* series, *Psychodopygus* Chagasi series, *Pressatia, Trichopygomyia* and *Trichophoromyia* should continue to be evaluated.

The sequencing of the *COI* gene allowed the detection of cryptic diversity within species. The species delimitation analysis of RESL and TCS splitted into at least two MOTUs the species: *Ps. panamensis, Mi. trinidadensis, Pi. evansi*,, and *Lu. gomezi*, which also achieved high rates of maximum intraspecific pairwise distances (Table 1) and were grouped in well-supported NJ clusters (Figure 2). In the first four cases, the detection of these genetic lineages may be associated with microevolutionary processes due to isolation by distance and geographic barriers (e.g., Andean region) since these four species presented clusters related to the geographic locations where they were sampled (Figure 2). This pattern has been seen in studies with a wide geographic distribution of the analyzed species [16, 19, 39, 54-55] or when clear geographic barriers are assessed, such as Amazonian riverbanks [14, 56] and caves in Thailand [18]. In the present study, a remarkable case comprising specimens of *Pi. evansi*, which has more than 8% of intraspecific genetic distance, were splitted into two MOTUs, the first comprising sequences from the Colombian department of Sucre and the other with sequences of specimens from Nicaragua and Honduras (Figures 1 and 2). These results raise the hypothesis that these populations represent distinct species, but the findings should be validated using integrative approaches to elucidate the actual taxonomic status of this species. Whether or not *Pi. evansi* represents different species, these molecular lineages may be taken into account from an epidemiological point of view because there may be variations regarding the ecological aspects and the vector-parasite interactions [57], and this should also be considered for the species *Ps. panamensis, Mi. trinidadensis*, and *Lu. gomezi* due to their vectorial role in transmitting *Leishmania* pathogens in the Neotropical region [3, 59].

The phylogenetic analysis of the *COI* gene allowed the formation of well-supported clades for the nominal species but failed to recover the evolutionary relationships of larger groups. Molecular markers of the mtDNA have a relatively high mutation rate, which is appropriate for identifying species and population structures but can fail in phylogenetic reconstructions of supraspecific relationships [59-60]. Indeed, the DNA barcoding approaches should not claim to establish evolutionary relationships based on a single rapidly evolving molecular marker, and other conserved genes such as rRNA 28S should be used for this purpose. Beyond that, even for conserved genes, multiple markers must be evaluated to generate species trees rather than gene trees [61]. However, some assumptions regarding *COI* barcode phylogenies can be raised when using appropriate methods of phylogenetic inference, such as Bayesian Inference and Maximum Likelihood. In the present work, some relationships between species were observed (Figure S1), and a well-supported clade was reconstructed for: i) all species of the *Lutzomyia* (*Tricholateralis*) subgenus; ii) two representatives of the Brumptomyiina subtribe and *Brumptomyia* genus; iii) the close-related species *Lu. lichyi* and *Lu. bifoliata* both of the *Lutzomyia* (*Lutzomyia*) subgenus; iv) *Pressatia camposi* and *Pr. choti*; v) two species of the *Psathyromyia* (*Forattiniella*) subgenus, *Pa. carpenter, Pa. aragaoi*; vi) *Trichophoromyia howardi* and *Th. velezbernali*; and vii) two well-supported clades within the *Nyssomyia* genus that are not necessarily related because of the low support value, the first containing the species *Ny. ylephiletor* and *Ny. trapidoi*, and the other comprising *Ny. fraihai, Ny. yuilli yuilli, Ny. yuilli pajoti, Ny. umbratilis*, and *Ny. antunesi*. In these last two cases, the clades were formed by the species that showed the lowest pairwise divergences in our dataset. There is no morphological evidence that the sampled *Nyssomyia* species are polyphyletic, so other approaches should assess the natural relationships between these taxa.

Furthermore, an interesting grouping pattern was observed for the Psychodopygina group since all species of this subtribe were grouped into a single clade, despite the low support value. The monophyly of the genus Psychodopygina appears to be consistent and has already been demonstrated in studies using multiple genetic markers [62] and analyzing a fragment of the rRNA 28S gene [63]. These efforts to confirm and establish the evolutionary relationships of sand flies using molecular data are hampered by the low coverage of sampled taxa.

## Conclusion

In summary, the sequencing and analysis of the *COI* DNA barcoding fragment allowed the correct delimitation of several Neotropical sand fly species from South and Central America. New important sequences of sand flies that had not been previously processed for this molecular marker were generated, which increases the relevance of DNA repositories so that the identification of sand flies is made more accurately using integrative tools. The findings of cryptic diversity within *Ps. panamensis, Mi. trinidadensis, Pi. evansi, Mi. cayennensis cayennensis*, and *Lu. gomezi* should be further evaluated to elucidate the possible presence of cryptic species, mainly considering the wide geographic distribution and the epidemiological importance of these species in transmitting pathogens to humans and other vertebrate hosts.

## Acknowledgments

We would like to offer our special thanks to the field team that made the collections, the PAHO for making the trip to the Central American countries possible, and the ecoepidemiology and Medical Entomology Units of PECET. We would also like to thank Andres Velez, Horacio Cadena, Rafael Vivero, and Luz Adriana Agudelo. Finally, we would like to thank PECET for funding this project at all stages and Dra Eunice Galati from FSP/USP Brazil for revising the manuscript.

## Author Contributions

**Conceptualization:** LPL, SIU

**Data Curation:** LPL, BLR

**Formal Analysis:** LPL, BLR

**Funding Acquisition:** IDV, SIU

**Investigation:** LPL, SIU

**Methodology:** LPL, BLR, SIU

**Resources:** IDV, SIU

**Visualization:** LPL, BLR

**Writing – Original Draft Preparation:** LPL, BLR

**Writing – Review & Editing:** LPL, BLR, IDV, SIU

